# Different Proteins Mediate Step-wise Chromosome Architectures in *Thermoplasma acidophilum* and *Pyrobaculum calidifontis*

**DOI:** 10.1101/2020.03.13.982959

**Authors:** Hugo Maruyama, Eloise I. Prieto, Takayuki Nambu, Chiho Mashimo, Kosuke Kashiwagi, Toshinori Okinaga, Haruyuki Atomi, Kunio Takeyasu

## Abstract

Archaeal species encode a variety of distinct lineage-specific chromosomal proteins. We have previously shown that in *Thermococcus kodakarensis*, histone, Alba, and TrmBL2 play distinct roles in chromosome organization. Although our understanding of individual archaeal chromosomal proteins has been advancing, how archaeal chromosomes are folded into higher-order structures and how they are regulated are largely unknown. Here, we investigated the primary and higher-order structures of archaeal chromosomes from different archaeal lineages. Atomic force microscopy of chromosome spreads out of *Thermoplasma acidophilum* and *Pyrobaculum calidifontis* cells revealed 10-nm fibers and 30–40-nm globular structures, suggesting the occurrence of higher-order chromosomal folding. Our results also indicated that chromosome compaction occurs toward the stationary phase. Micrococcal nuclease digestion indicated that fundamental structural units of the chromosome exist in *T. acidophilum* and *T. kodakarensis* but not in *P. calidifontis* or *Sulfolobus solfataricus*. In vitro reconstitution showed that, in *T. acidophilum,* the bacterial HU protein homolog HTa formed a 6-nm fiber by wrapping DNA, and that Alba was responsible for the formation of the 10-nm fiber by binding along the DNA without wrapping. Remarkably, Alba could form different higher-order complexes with histone or HTa on DNA in vitro. Mass spectrometry detected HTa in the *T. acidophilum* chromosome but not in other species. A putative transcriptional regulator of the AsnC/Lrp family (Pcal_1183) was detected on the *P. calidifontis* chromosome, but not on that of other species studied. Putative membrane-associated proteins were detected in the chromosomes of the three archaeal species studied, including *T. acidophilum*, *P. calidifontis*, and *T. kodakarensis*. Collectively, our data show that Archaea use different combinations of proteins to achieve chromosomal architecture and functional regulation.

## 1. Introduction

Genomic DNA needs to be folded properly in the cell to simultaneously accomplish compaction of the genetic material and accessibility to the transcription and replication machinery in all three domains of life. In eukaryotic cells, genomic DNA folding is primarily achieved through the nucleosome, a structure composed of histone octamers with approximately 150 bp of DNA wrapped around them (Luger et al., 1997; Davey et al., 2002). Additional proteins, including linker histone H1 and DNA topoisomerases, have been proposed to contribute to the higher-order folding of the eukaryotic genome (Thomas, 1984; Woodcock et al., 2006; Hizume et al., 2007; Fyodorov et al., 2018). Chromosome structural regulation is also pivotal for cell cycle progression. For example, condensin, a structural maintenance of chromosomes (SMC)-superfamily protein, is required for mitotic chromosome condensation in eukaryotes (Hirano et al., 1997; Ono et al., 2003). Epigenetic modifications also reportedly regulate gene expression (Grewal and Moazed, 2003; Li and Reinberg, 2011). Moreover, recent technological advances in chromosome conformation capture (3C) have shown that the eukaryotic genome is hierarchically folded into chromosomal domains, such as topologically associating domains (TADs) (Gibcus and Dekker, 2013; Barutcu et al., 2017). In bacteria, genome compaction is achieved through the functions of nucleoid-associated proteins (NAPs) including HU, Fis, H-NS, and IHF (Skoko et al., 2006; Sarkar et al., 2009; Wang et al., 2013; Badrinarayanan et al., 2015; Hammel et al., 2016; Japaridze et al., 2017). Non-protein factors, such as macromolecular crowding, also contribute to the higher-order folding and regulation of the bacterial nucleoid (De Vries, 2010). It has also become evident that the bacterial nucleoid is segmented into many highly self-interacting regions called chromosomal interaction domains (CIDs), which are equivalent to the eukaryotic TADs, and the CIDs are further organized into large spatially distinct domains called macrodomains (Dame and Tark-Dame, 2016; Brocken et al., 2018; Verma et al., 2019).

Archaea constitute one of the three domains of life, along with Eukarya and Bacteria. Although Archaea are considered extremophiles that live in extreme environments, including high temperature, high salinity, or low pH (Forterre, 2016), they have also been found in moderate environments, including marine environments (Lloyd et al., 2013), soil (Bates et al., 2011), and even the human body (Probst et al., 2013; Koskinen et al., 2017; Nkamga et al., 2017; Pausan et al., 2019). Moreover, although no archaeal pathogens have been identified, Archaea have been associated with human diseases, including periodontal disease (Lepp et al., 2004) and inflammatory bowel disease (Blais Lecours et al., 2014). Thus, understanding how Archaea respond to and influence their environment has become crucial. Whereas control of gene expression in Archaea is achieved in part through transcription factors (TFs) resembling bacterial TFs, recent studies have suggested that chromosome structure and the interplay between chromosomal proteins and basal transcription machinery might also play important roles in gene expression (Peeters et al., 2015; Lemmens et al., 2019).

Archaeal chromosomal proteins are diverse (Sandman and Reeve, 2005; Luijsterburg et al., 2008; Driessen and Dame, 2011). For example, most species in Euryarchaeota, one of the major archaeal phyla, encode proteins homologous to eukaryotic histone (Nishida and Oshima, 2017; Henneman et al., 2018). Species in newly proposed phyla such as Nanoarchaeota, Thaumarchaeota, and Lokiarchaeota also encode histone (Henneman and Dame, 2015; Henneman et al., 2018). Exceptions in Euryarchaeota are the members of the order *Thermoplasmatales,* which lack histones and instead encode HTa (**H**istone-like protein of ***T**hermoplasma **a**cidophilum*), a protein homologous to bacterial HU (DeLange et al., 1981; Hocher et al., 2019). Whether HTa has a role similar to that of bacterial HU or an Archaea-specific function remains elusive. Proteins such as Cren7, CC1, and Sul7 are specific to species in Crenarchaeota, another major archaeal phylum (Driessen and Dame, 2011). Alba, a 10-kDa DNA/RNA-binding protein, is found in both Euryarchaeota and Crenarchaeota (Guo et al., 2003; Laurens et al., 2012), as well as in newly proposed phyla including Nanoarchaeota, Korarchaeota, Thaumarchaeota, and Lokiarchaeota (Henneman and Dame, 2015; Sanders et al., 2019). Alba has been shown to undergo post-translational modifications including methylation and acetylation (Bell et al., 2002; Cao et al., 2018). Alba has also been shown to have several different modes of interaction with DNA, such as DNA stiffening or bridging (Laurens et al., 2012). However, how Alba cooperates with other chromosomal proteins in higher-order chromosome folding and whether it plays different roles in Euryarchaeota and Crenarchaeota remain to be elucidated. Besides these DNA-binding proteins that are usually smaller than 10 kDa, some larger transcription factor-like proteins also behave as chromosomal proteins in Archaea. TrmB-like 2 (TrmBL2), a protein homologous to the sugar-responsive transcriptional regulator TrmB, is an abundant chromosomal protein that is completely conserved among *Thermococcales* (Gindner et al., 2014; Kim et al., 2016). Lines of experimental evidence suggest that TrmBL2 is a chromosome-associated protein resembling bacterial H-NS, with functions that include filament formation on DNA and the global suppression of gene expression (Maruyama et al., 2011; Efremov et al., 2015). More recently, Hi-C chromosome conformation capture experiments on *Sulfolobus* species showed that a new class of SMC protein coalescin mediates the crenarchaeal chromosome into a two-domain organization resembling eukaryotic A/B compartments (Takemata et al., 2019).

In the present study, we analyzed the protein composition and structure of chromosomes from diverse archaeal lineages, aiming to understand the general principles of genome folding. Species used in this study belong to either Euryarchaeota or Crenarchaeota, two major phyla in Archaea, and encode different combinations of chromosomal proteins. *Thermococcus kodakarensis* is a euryarchaeon that encodes histone, Alba, and TrmBL2 as major chromosomal proteins (Maruyama et al., 2011). *Thermoplasma acidophilum* is a euryarchaeon that is an exception, as it lacks histones, but it encodes a protein HTa homologous to bacterial HU, together with Alba (Ruepp et al., 2000). Since it is unclear whether HTa plays roles similar to that of bacterial HU, we investigated how HTa contributes to genome folding. *Pyrobaculum calidifontis* is a hyperthermophilic crenarchaeon that encodes Alba, Cren7, CC1, and Sso10a (Amo et al., 2002; Peeters et al., 2015). *Sulfolobus solfataricus* is a crenarchaeon that encodes Alba, Cren7, Sul7, and Sso10a (Peeters et al., 2015). Both the primary and higher-order chromosome structures were analyzed with atomic force microscopy (AFM). Chromosomal proteins in each species were identified through a combination of chromosome isolation and mass spectrometry. The minimal structural unit of each chromosome was investigated with micrococcal nuclease (MNase) digestion analysis and the in vitro reconstitution of protein-DNA complexes.

## 2. Results

### 2.1. The archaeal chromosome is commonly composed of 10-nm fibers and 30-40-nm globular structures

We previously showed that, in *T. kodakarensis*, the chromosome is more compact in the stationary phase than in the log-phase (Maruyama et al., 2011). Here, the extent of chromosome compaction in two archaeal species from different phyla, *T. acidophilum* and *P. calidifontis*, were analyzed by using the on-substrate lysis method (see Materials and Methods). Cells were grown and applied on coverslips, then mildly lysed with detergent. Staining of the lysed cells with 4’,6-diamidino-2-phenylindole (DAPI) showed stretched and decondensed chromosome released from log-phase cells for both archaeal species (Figure 1A). In contrast, compact chromosome structures were dominant after the lysis of stationary phase cells based on DAPI staining (Figure 1B). These results suggest the difference in chromosome organization of archaeal cells between the growth phases.

**Figure 1.**
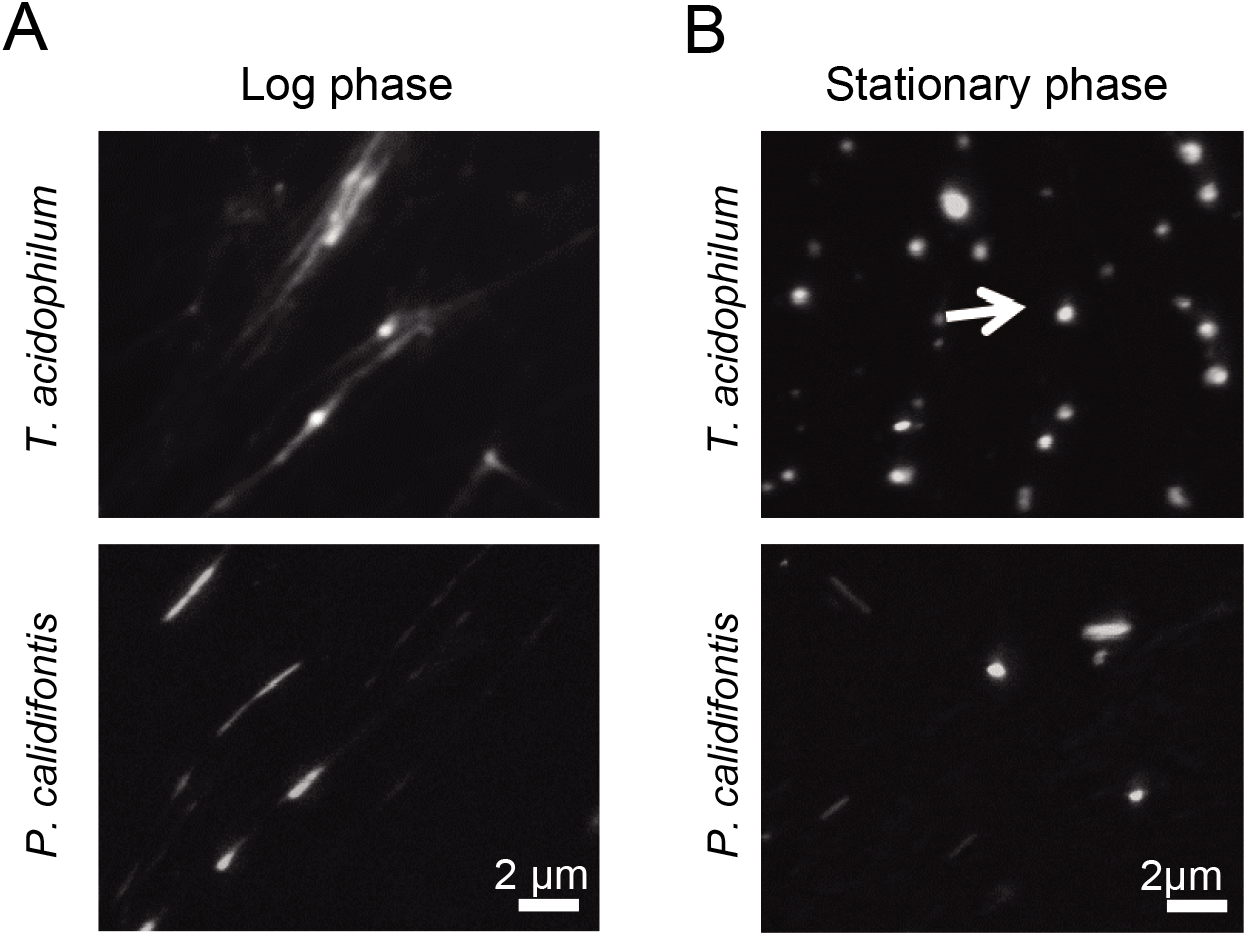
Genomic material released from *Thermoplasma acidophilum* and *Pyrobaculum calidifontis* cells. DAPI staining was performed after on-substrate lysis of the cells during (A) log and (B) stationary phases. Fluorescence microscopy images are shown. (A) Chromosomes were decondensed and elongated in the log-phase for both *T. acidophilum* (upper panel) and *P. calidifontis* (lower panel). (B) The stationary phase chromosome was more compact than the logphase chromosome (arrow). Scale bars: 2 μm.

Detailed AFM analysis of chromosomes released from the lysed cells identified two distinct chromosomal structures common to both species, 10-nm fiber and 30–40-nm globular structures. The 10-nm fiber structure was prevalent on the well-spread chromosome in the log-phase in both species (Figure 2A). The diameter of the fibers was 10.1 ± 4.2 nm (mean ± SD, n = 119) in *T. acidophilum* and 11.5 ± 4.2 nm (n = 110) in *P. calidifontis* (Figure 2B). Larger globular structures were found where chromosomes were less spread in both species in the stationary phase, wherein most cells showed limited chromosome release (Figure 2C). The diameters of the globular structures differed between the two species and were specifically 38.6 ± 10.1 nm (n = 217) nm in *T. acidophilum* and 29.1 ± 5.1 nm (n = 152) nm in *P. calidifontis* (Figure 2D), suggesting differences in the fundamental structure or the higher-order architecture of the chromosome. Although the fiber and globular structures are seen in both growth phases, these results show that the extent of archaeal chromosome folding is greater in the stationary phase.

**Figure 2.**
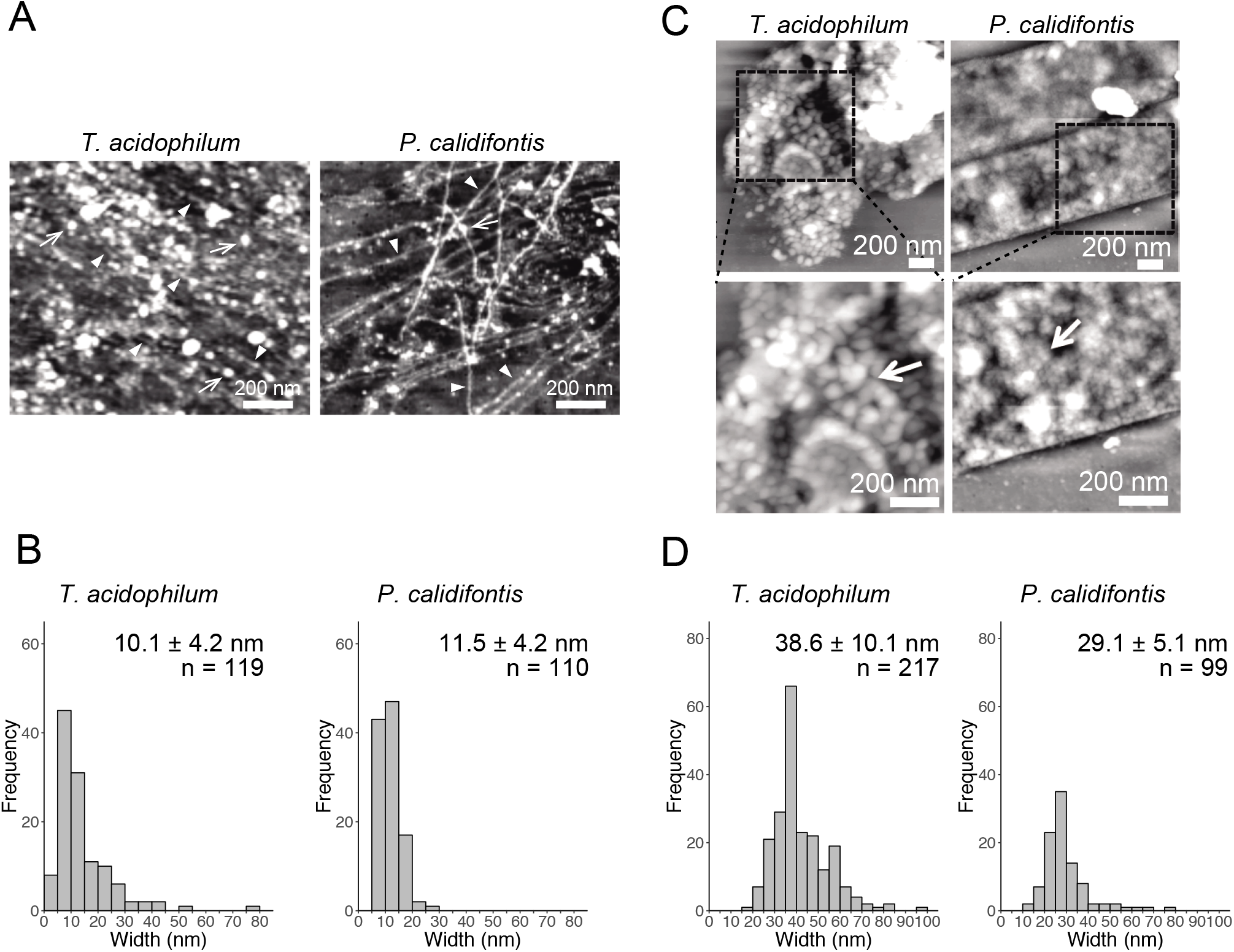
Structural details of genomic materials from *Thermoplasma acidophilum* and *Pyrobaculum calidifontis* after on-substrate lysis visualized with atomic force microscopy (AFM). (A-B) Extensively released chromosome fibers of log-phase cells. (A) AFM images of the released genomic materials show relatively thin fibers (filled triangles) and globular structures (arrows). (B) Histograms indicate the diameters of the released fiber structures indicated by filled triangles in (A). (C-D) Compact chromosome of stationary phase cells. (C) AFM images of the lysed cells. Boxed areas in the upper panels were shown in detail in the lower panels. Globular structures (arrows) were observed more frequently than in the log phase (D) Histograms indicate the diameters of the globular structures. Scale bars: 200 nm.

### 2.2. Minimal unit of the archaeal chromosome revealed with MNase digestion

On-substrate lysis results revealed that the size of the primary structure of the archaeal chromosome is approximately 10 nm in diameter (Figure 2A and 2B). We next investigated the minimal structural unit through MNase digestion of purified chromosomes from four archaeal species: *T. kodakarensis* and *T. acidophilum* from Euryarchaeota, and *P. calidifontis* and *S. solfataricus* from Crenarchaeota. The MNase digestion was performed without chemical fixation. The analysis revealed a variation in minimal structural units in each species (Figure 3). As we have previously shown, MNase digestion of the *T. kodakarensis* chromosome resulted in a 30-bp laddered pattern with a minimum of 60 bp of DNA (Figure 3A). The 30-bp intervals detected are typical of some histone-containing Archaea and reflect the flexible histone multimeric structure, which was shown previously (Maruyama et al., 2013; Mattiroli et al., 2017; Henneman et al., 2018). Interestingly, MNase digestion of the *T. acidophilum* chromosome resulted in the accumulation of 40~50 bp or 80~90 bp of DNA depending on the enzyme concentration (Figure 3B), corresponding with the results of the previous reports (Searcy and Stein, 1980; Hocher et al., 2019). The 40~50 bp fragment suggests the size of DNA involved in the primary unit. The larger DNA fragment of 80~90 bp identified with a lower MNase concentration indicated that this unit can be positioned adjacent to each other (Hocher et al., 2019). In contrast, the MNase digestion of the chromosomes of the two crenarchaeal species, *P. calidifontis* and *S. solfataricus*, did not yield the accumulation of DNA fragments of a particular size (Figure 3C and 3D). These results demonstrate that in crenarchaeal species, which encode neither histone nor HTa, the chromosomes do not have a defined-size structural unit that protects the genomic DNA from MNase digestion. Whereas on-substrate cell lysis revealed similar fibrous chromosome structures among *T. acidophilum* and *P. calidifontis* (Figure 2), as well as in *T. kodakarensis*, as previously reported (Maruyama et al., 2011), the MNase digestion patterns indicated that the fine details in the organization of these structures vary, probably due to differences in the chromosomal proteins involved.

**Figure 3.**
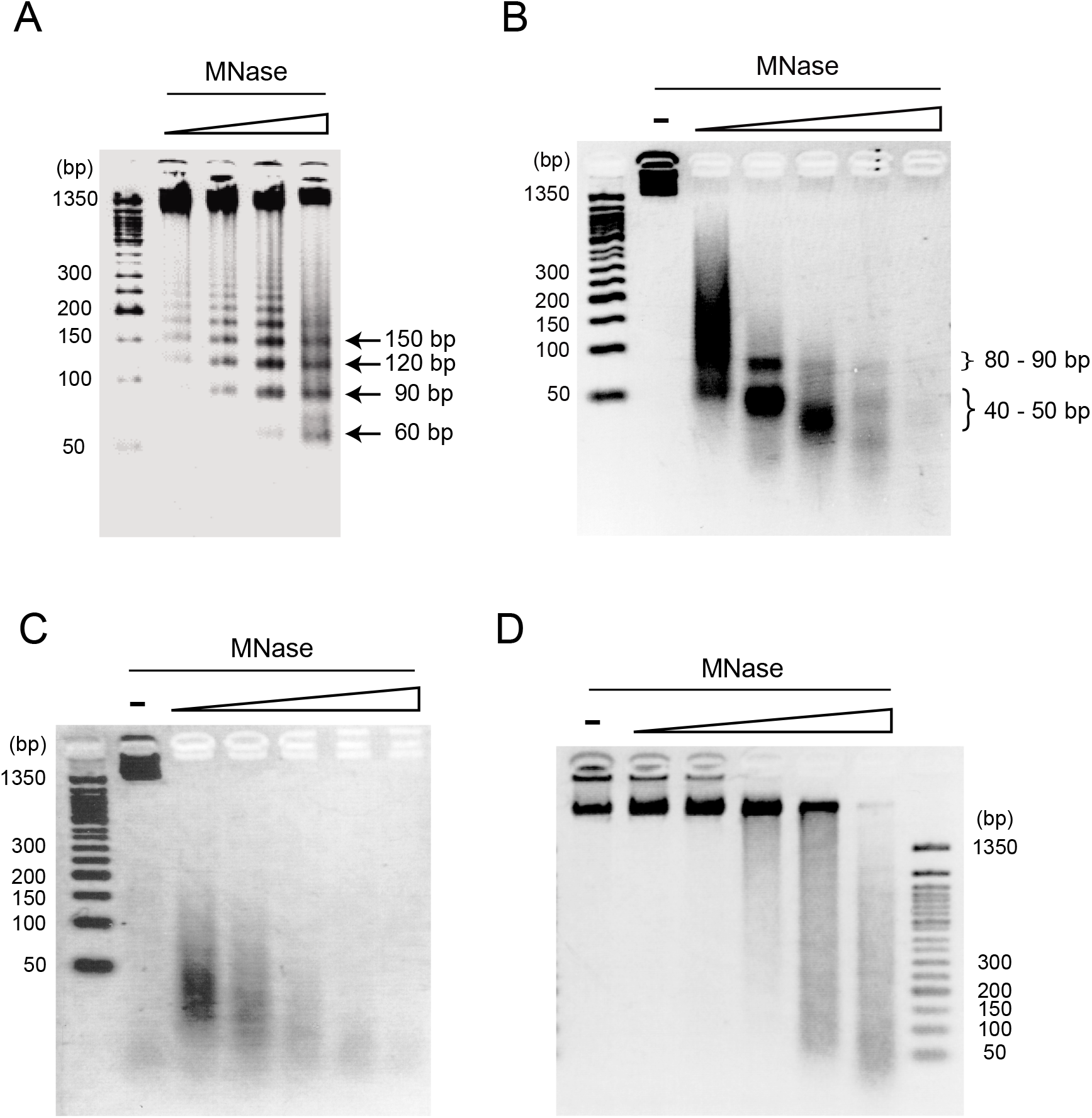
Digestion pattern of archaeal chromosomes treated with micrococcal nuclease (MNase). Purified chromosomes of (A) *Thermococcus kodakarensis*, (B) *Thermoplasma acidophilum*, (C) *Pyrobaculum calidifontis*, and (D) *Sulfolobus solfataricus* were digested with increasing concentrations of MNase and separated on 2.5% agarose gels in 1X TBE. The accumulation of DNA of particular sizes was observed with *T. kodakarensis* (arrows) and *T. acidophilum* (curly brackets) but not with *P. calidifontis* and *S. solfataricus* chromosomes. MNase concentration was 0.3, 1, 3, 10 U MNase in 100 μl reaction (A) or 0, 0.3, 1, 3, 10 and 30 U MNase in 100 μl reaction (B-D).

### 2.3. Mass spectrometry identifies chromosome-associated proteins in archaeal species

Protein components of isolated chromosomes of *T. acidophilum*, *P. calidifontis* and *T. kodakarensis* were analyzed using mass spectrometry (Figure 4). Tables 1 to 3 provide an overview of the distribution of chromosomal proteins in each archaeal species. Proteins predicted to be involved in replication, transcription, and translation were detected in the three species. These include the DNA primase DnaG detected in *T. acidophilum* and *P. calidifontis* and DNA-directed RNA polymerase subunits detected in *T. acidophilum* and *T. kodakarensis*. Ribosome subunits were commonly found in all species and constituted a large portion of the identified proteins, which is reasonable considering that transcription and translation are coupled in Archaea (French et al., 2007).

**Figure 4.**
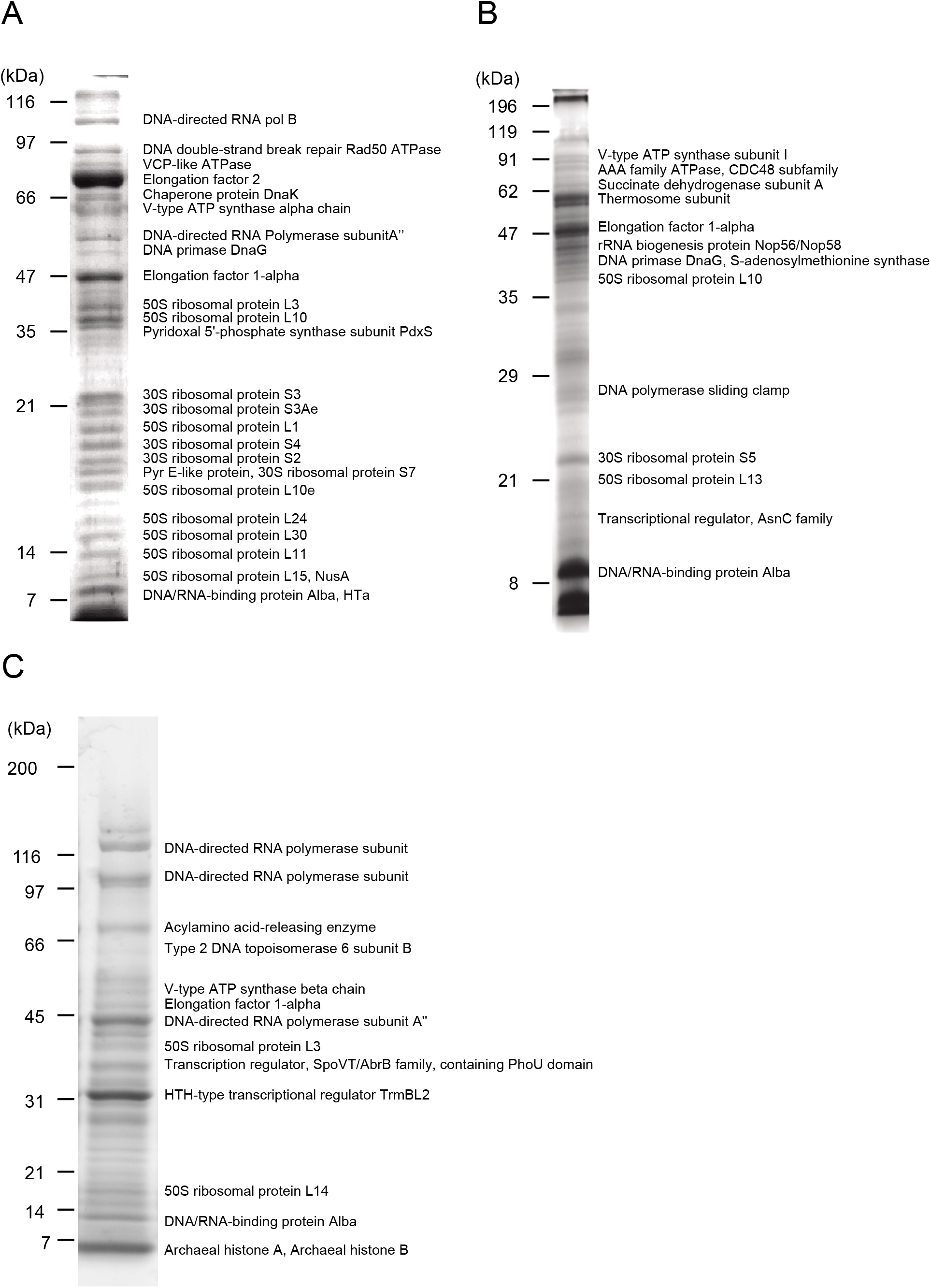
Protein components of isolated chromosomes. Chromosomes Isolated fron (A) *T. acidophilum* (B) *P.calidifontis* (C) *T. kodakarensis* were digested with MNase, separated through SDS-PAGE and stained with Comassie Brilliant Blue. Representative proteins detected with mass spectrometry are indicated next to the corresponding bands. See Tables 1 – 3 for details.

Apart from these general information processing proteins, DNA-binding proteins that might contribute to chromosome folding and regulation were also detected. Alba was detected in all three species studied (Figure 4 and Tables 1–3). In the *T. acidophilum* chromosome, HTa was detected together with Alba (Figure 4A and Table 1). In the *P. calidifontis* chromosome, a protein annotated as ‘Transcriptional regulator, AsnC family’ (tremble ID: A3MVE1, gene ID: Pcal_1183) was detected in addition to Alba (Figure 4B and Table 2). It has been suggested that some AsnC/Lrp family transcriptional regulators have global gene regulatory roles as well as an architectural function, instead of being a specific transcriptional regulator (Peeters et al., 2015). The identification of Pcal_1183 in *P. calidifontis,* which has not been reported as a chromosomal protein, supports the view that TF-like proteins act as chromosome proteins in Archaea, as is the case for TrmBL2 in *Thermococcales* (Maruyama et al., 2011). Cren7, although encoded in the *P. calidifontis* genome, was not detected in the present mass spectrometry analysis, suggesting that the amount of Cren7 on the *P. calidifontis* chromosome is lower than the detection limit of our method. In *T. kodakarensis*, histones, Alba, and TrmBL2 were detected as previously reported (Figure 4C and Table 3) (Maruyama et al., 2011).

Interestingly, in addition to proteins with apparent DNA-binding ability, a few putative membrane-associated proteins were detected in the chromosome. For example, V-type ATPases were found in all three species (Figure 4 and Tables 1–3), suggesting a general role of these proteins in archaeal chromosome folding and regulation. Because the chromosome isolation procedure included cell lysis with a detergent and sucrose density gradient sedimentation, we assume that the isolated chromosome was not contaminated with cell membrane or membrane-bound proteins, as shown previously for *T. kodakaresis* (Maruyama et al., 2011).

### 2.4. Distinct fundamental structures formed with Alba and HTa

Alba was common among the three species studied (Figure 4 and Tables 1–3). When recombinant Alba proteins from *T. acidophilum* or *P. calidifontis* genes expressed in *E. coli* were mixed with linear 3-kb DNA in vitro, a fiber structure was formed at Alba:DNA weight ratio higher than 6:1 (Figure 5A). The diameter of the fiber was 10.0 ± 2.8 nm (n = 80) in *T. acidophilum* and 12.1 ± 2.2 nm (n = 119) in *P. calidifontis* (Figure 5B). These correlate well with the structures visualized in the on-substrate lysis experiment (Figure 2A and 2B), suggesting that Alba is responsible for the formation and maintenance of the 10-nm fiber structures. The contour length of the filamentous structures was not shorter compared to the theoretical length of a 3-kb DNA, which is ~1000 nm (Maruyama et al., 2011), indicating that DNA does not wrap around Alba molecules (Figure 5C). At relatively higher Alba concentrations, these filaments seemed to be capable of folding onto themselves and forming more complex structures. The results strongly support a role of Alba protein contributing to primary order folding of the DNA into 10-nm fibers. In addition, the data suggest the role of Alba in forming more complex, higher-order structures.

**Figure 5.**
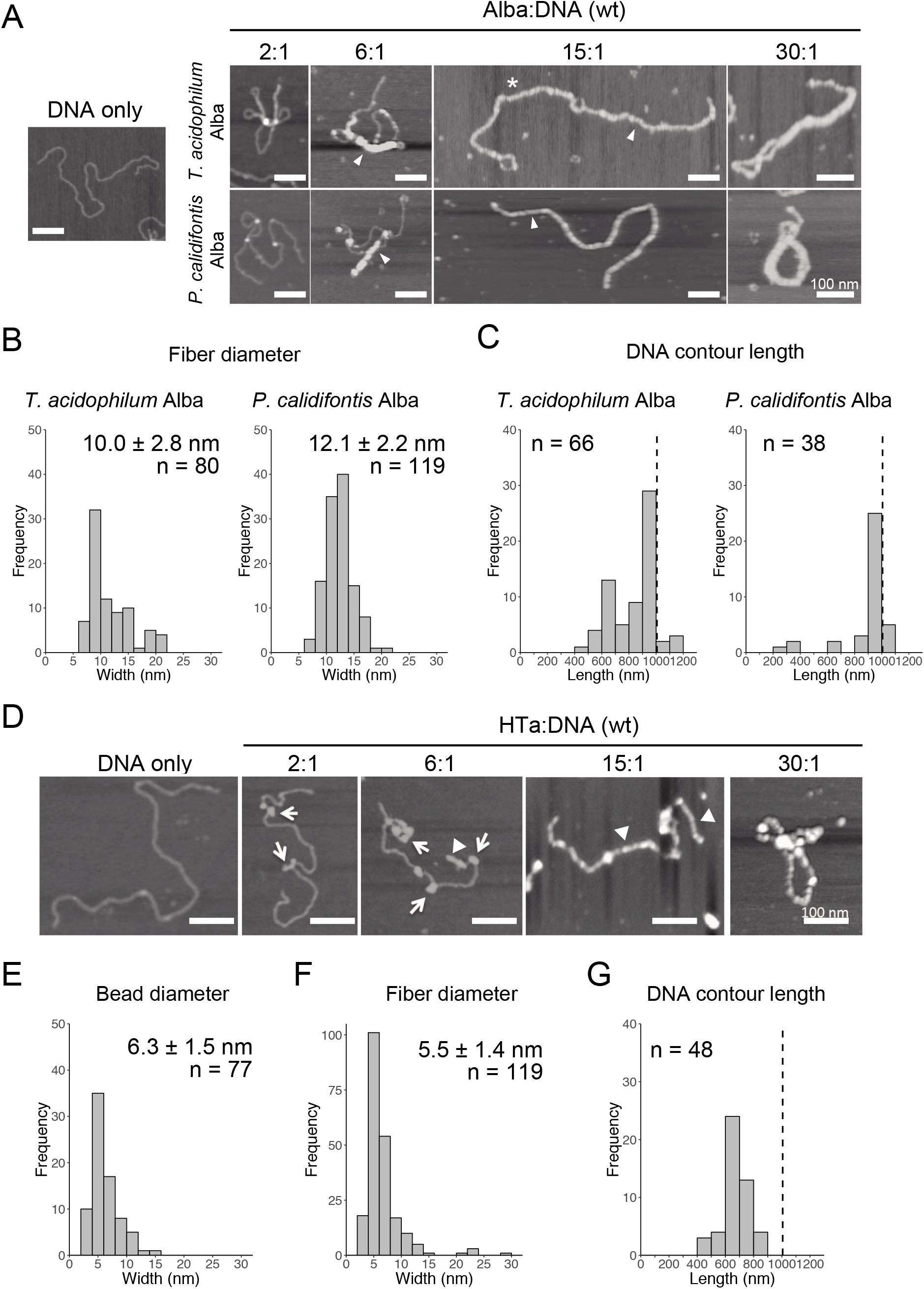
In vitro reconstitution of chromosome structures with recombinant Alba and HTa. (A-C) Reconstitution on a linear 3-kb plasmid using recombinant Alba from *Thermoplasma acidophilum* and *Pyrobaculum calidifontis* at different Alba to DNA weight ratios. (A) AFM images showing various structures formed with Alba and DNA. Fibers are indicated with triangles. Alba:DNA weight ratios are indicated. The asterisk indicates a structure in which two 3-kb DNA molecules are joined together by Alba binding. (B) Histograms show the diameters of the fibrous structures formed (indicated with triangles in (A)). (C) Histograms show the respective contour lengths of DNA at a 15:1 Alba:DNA ratio. The theoretical length of a linear 3-kb DNA (~1000 nm) is indicated with a dashed line. See Figure 5A for a histogram of unbound DNA length. (D-G) Reconstitution with histidine-tagged HTa from *T. acidophilum* at varying protein to DNA ratio. (D) AFM images showing beaded (arrows) and filamentous (triangles) structures. (E) Diameter of the beads formed at a relatively lower HTa concentration. (F) Width of the filaments formed at relatively higher HTa concentration. (G) Histogram shows the contour DNA lengths of the structures formed at a 15:1 HTa:DNA ratio. Dashed line indicates the theoretical length of a 3-kb unbound DNA. Scale bars: 100 nm.

Whereas most Euryarchaeota commonly encode histones, species belonging to *Thermoplasmatales* lack histones and encode HTa, a homolog of bacterial HU. It has been proposed that *Thermoplasmatales* acquired the HU gene from bacteria through horizontal gene transfer (HGT) and lost the histone gene and that HTa plays a role similar to that of archaeal histone (Hocher et al., 2019). We investigated how HTa interacts with DNA at the primary level. The reconstitution using recombinant histidine-tagged HTa from *T. acidophilum* showed a beaded structure (Figure 5D), similar to the beads-on-a-string structure formed with archaeal histone (Maruyama et al., 2011; Maruyama et al., 2013) or bacterial HU (Rouvière-Yaniv et al., 1979). The diameter of the HTa beads was 6.3 ± 1.5 nm (n = 77) (Figure 5E), which is thinner than the 10-nm Alba filaments (Figure 5A and 5B). At higher concentrations, HTa formed a filament of 5.5 ± 1.4 nm in width (n = 221) on the DNA (Figures 5D and 5F). The contour DNA length of reconstituted nucleoprotein structure was shorter than that of naked DNA (Figure 5G), reflecting DNA wrapping around HTa particles. These results support the notion that archaeal HTa is functionally more closely related to histones than bacterial HU (Hocher et al., 2019). The ~6-nm particle might be a fundamental unit of the *Thermococcales* chromosome, together with the 10- nm fiber formed with Alba (Figure 5A and 5B).

### 2.5. Relationship between Alba and other chromosomal proteins in forming structural complexes with DNA

Alba can form various structures with DNA, depending on its amount (Figure 5A). This flexibility might facilitate its interaction with other structural proteins, forming DNA-based complexes. Alba coexists with other DNA-binding proteins in the cell. For example, *T. kodakarensis* encodes histone and Alba, whereas *T. acidophilum* encodes HTa and Alba. We therefore combined Alba with the chromosomal proteins, histone and HTa, to determine how they cooperate in compacting DNA.

In vitro reconstitution using histone alone at a 1:1 histone:DNA weight ratio resulted in compact beaded fiber structures displaying decreased contour length of the DNA-protein complex (Figure 6A). Next, histone and Alba were mixed together with DNA. Alba was added in the concentration range at which it started to form the 10-mn fiber structure when it was mixed with DNA alone (Figure 5A). Reconstitution using both histone and Alba from *T. kodakarensis* at a 1:3 or 1:7 histone:Alba ratio resulted in the formation of structures somewhat similar to those formed with DNA and histone alone, (Figure 6A), but with a larger proportion of DNA molecules having a longer contour length (> 500 nm) (Figure 6A). As Alba does not wrap DNA (Figure 5A and 5B), this result suggests that Alba binding decreases the extent of histone-mediated DNA folding or that Alba competes with histone for DNA-binding, leading to less compact structures. Although the amount of Alba relative to histone in *T. kodakarensis* cell (3:1 histone:Alba in weight) (Maruyama et al., 2011) is estimated to be lower than that used in this experiment, these results indicate that Alba can compete with histone at least at the local level.

**Figure 6.**
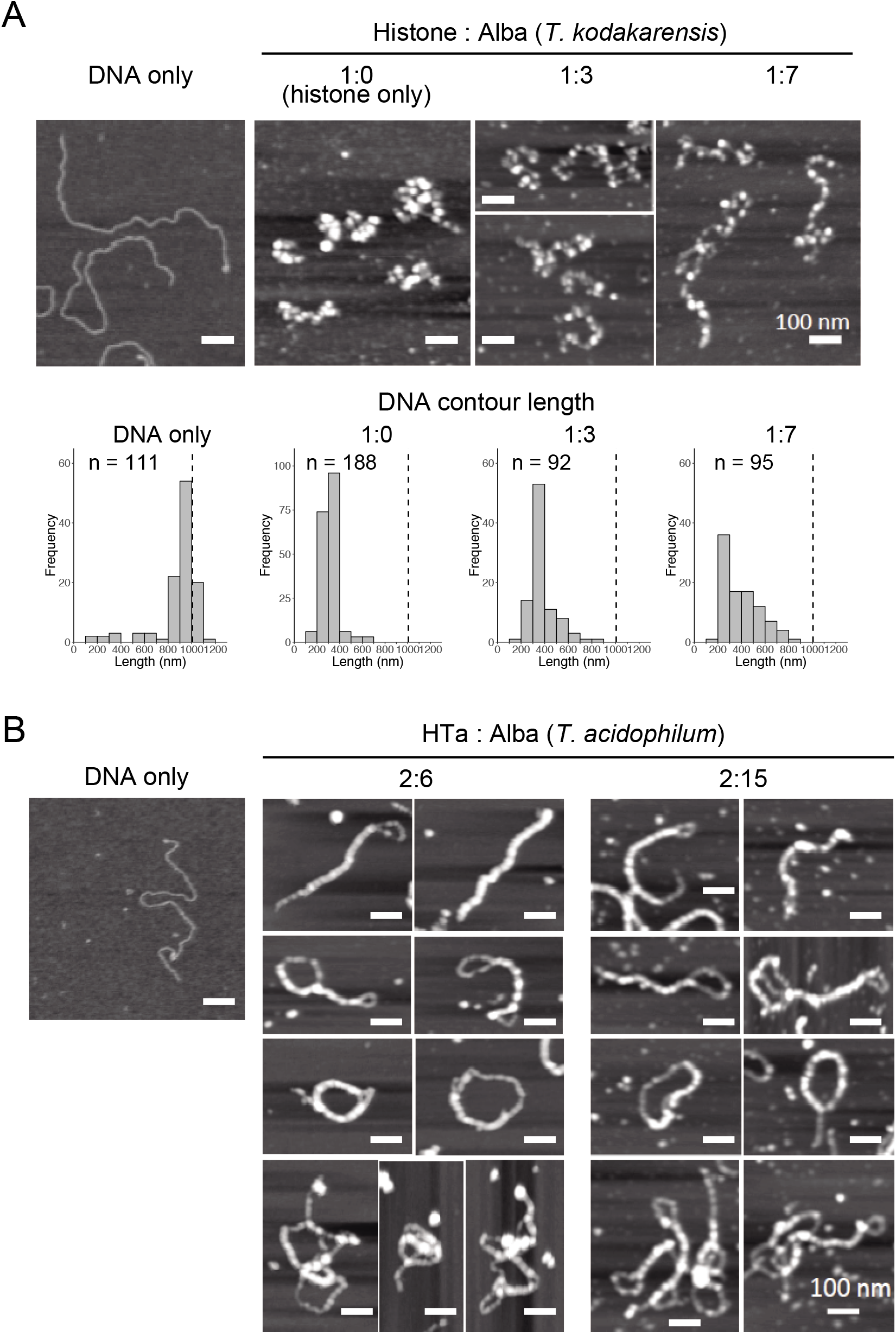
In vitro reconstitution using a combination of archaeal proteins. Chromatin structures were reconstituted on a linear 3-kb plasmid using recombinant proteins at varying Alba concentrations. (A) AFM images showing reconstitution using histone and Alba from *Thermococcus kodakarensis*. Histograms below the AFM images show the contour DNA length in each condition. Dashed line indicates the theoretical length of a 3-kb unbound DNA. (B) AFM images showing reconstitution using HTa and Alba from *Thermoplasma acidophilum.* Scale bars: 100 nm. Note that histograms are not shown because the correct contour DNA length were unable to measure due to a high degree of folding or joining of the fiber structures.

Next, HTa and Alba from *T. acidophilum* were mixed together with DNA (Figure 6B). The concentration of HTa was fixed at which it formed separate beaded structure on DNA (1:2 DNA:HTa) (Figure 5D). Alba was mixed at the concentration range at which it formed fiber structure on DNA (1:6 or 1:15 DNA:Alba) when it was mixed alone with DNA (Figure 5 A). Reconstitution using HTa and Alba from *T. acidophilum* at 2:6 and 2:15 weight ratios (Figure 6B) resulted in structures similar to those formed by Alba alone (Figure 5A). Structures with DNA strands joined together were dominant, and DNA molecules were considerably folded (Figure 6B). Moreover, some complexes were able to fold upon themselves, providing additional compaction (Figure 6B). These data indicate that unlike with histone (Figure 6A), Alba with HTa facilitates DNA compaction through DNA strand bridging (Figure 6B).

Thus, Alba encoded in each species might play different roles depending on the interaction or synergy with other proteins associated with the chromosome in each archaeal lineage. Binding of HTa might increase the efficacy of Alba-mediated DNA bridging, leading to complexes that are more compact than those with either HTa or Alba alone. Whether this compaction is achieved through local structural change in chromatin due to the binding of individual protein or through direct protein-protein interaction between HTa and Alba needs to be elucidated.

## 3. Discussion

In the present study, we investigated the fundamental and higher-order structures of archaeal chromosomes from different lineages. We show that archaeal chromosomes comprise step-wise structures that consist of primary fibrous structures of 10 nm in diameter and higher-order globular structures of 30–40 nm. The minimal unit of the chromosome varies among species. Archaeal chromosomes undergo compaction toward the stationary phase (Figure 7A). We also show that archaeal HTa can wrap DNA and form a 6-nm beaded fiber, which might be folded into the higher-order structures in association with Alba in *T. acidophilum*. Interestingly, TF-like proteins were found in *T. kodakarensis* and *P. calidifontis*. Membrane-associated proteins were detected in the chromosomes of all three species. Finally, we demonstrate that Alba-mediated chromosomal structures vary depending on the presence and identity of additional architectural proteins.

**Figure 7.**
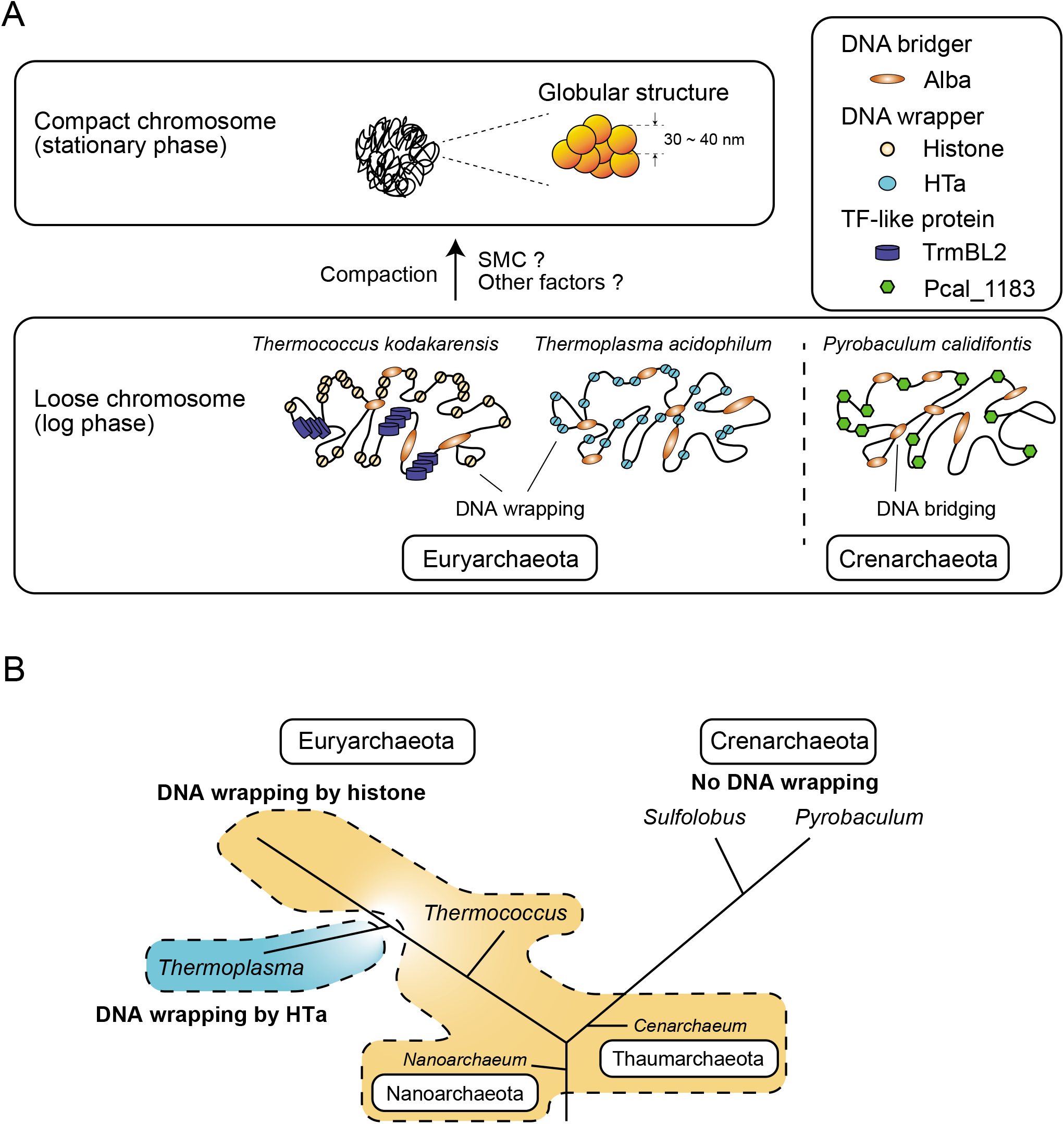
Model of archaeal chromosome folding. (A) In Euryarchaeota, histone or HTa wraps genomic DNA, whereas there is no DNA wrapper in Crenarchaeota. Alba can bridge distant genomic DNA. Chromosome compaction occurs toward the stationary phase, when 30~40-nm globular structures are formed. (B) Schematic phylogenetic tree of selected Archaea (Sandman and Reeve, 2005) together with the distributions of histone and HTa. Species in Euryarchaeota utilize histone or HTa *(Thermoplasmatales)* to wrap DNA at the primary level. Members of Nanoarchaeota and Thaumarchaeota also encode histone that might wrap DNA. Species in Crenarchaeota neither encode histone nor wrap DNA. The branch lengths do not correspond to evolutionary distances. Note that *Cenarchaeum* used to be classified as a member of Crenarchaeota, but is now considered a member of Thaumarchaeota (Spang et al., 2010).

### 3.1 Primary and higher-order folding of archaeal chromosomes

Higher-order chromosome architecture and its regulation have been extensively studied in bacteria and eukaryotes. In bacteria, the coordination of various NAPs is required for proper folding and regulation of the nucleoid (Verma et al., 2019). For example, the *Escherichia coli* nucleoid is composed of 10-, 40-, and 80-nm fibers and undergoes compaction in the stationary phase, in which Dps is highly expressed (Ali Azam et al., 1999; Kim et al., 2004). Oxidative stress also causes Dps-dependent nucleoid compaction in *Staphylococcus aureus* (Morikawa et al., 2019). In eukaryotes, it has been viewed that beads-on-a-string chromatin is folded into a 30- nm fiber with the linker histone H1 (Hizume et al., 2005; Prieto and Maeshima, 2019). Our results indicate that archaeal chromosomes also comprise step-wise structures, sharing features with both bacterial and eukaryotic chromosomes. A hierarchical mode of archaeal genome packing can be inferred wherein the 6- or 10-nm fiber structures form the 30–40-nm globular structures. The 6- or 10-nm fibers might represent the first order of folding in the archaeal chromosome, wherein DNA is bound and constrained by one or more abundant chromosomal protein(s). The abundant chromosomal proteins present in each species and their interactions with each other might contribute to species-specific primary and higher-order chromosome structures.

We show that the archaeal chromosome undergoes compaction toward the stationary phase, similarly to what happens in the bacterial nucleoid. However, how compaction is achieved remains elusive. SMC-superfamily proteins are known to be involved in higher-order architecture and the regulation of chromosomes in eukaryotes and bacteria (Uhlmann, 2016). The Hi-C experiments on *Sulfolobus* species showed that an SMC-family protein coalescin contributes to the formation of distinct chromosome domains that undergo descrete and specific higher-order interactions (Takemata et al., 2019). Although SMC homologs are present in archaeal genomes based on genomic sequences, they were not detected using mass spectrometry. Low amounts of SMC proteins might be sufficient for their function in archaeal cells, probably cooperating with other chromosomal proteins to achieve higher-order folding of the chromosome (Figure 7A). An indirect association of SMC protein with the chromosomes might also be a reason why they were not detected in our mass spectrometry analysis. Indeed, bacterial SMC proteins are loaded onto the chromosome by ParB proteins (Marbouty et al., 2015; Wang et al., 2015). Post-translational modifications of archaeal DNA-binding proteins such as Alba and Cren7 have been reported (Bell et al., 2002; Niu et al., 2013; Cao et al., 2018). Detailed analyses of the influence of such post-translational modifications on protein-protein and protein-DNA interactions might also provide insights into mechanisms of chromosome folding and regulation.

### 3.2. Role of HTa in *T. acidophilum*

HTa is an abundant protein in members of *Thermococcales*, which lack histones. Our present study shows that *T. acidophilum* HTa can form a beads-on-a-string structure by wrapping DNA and that the wrapping protects DNA from MNase digestion (Figure 3B and 5D). Taken together with a recently reported analysis of HTa (Hocher et al., 2019), these results indicate that HTa has characteristics more similar to those of archaeal histone than to those of bacterial HU. Moreover, HTa was recently reported to form oligomers (Hocher et al., 2019) resembling the flexible “hypernucleosome” formed by multiple archaeal histone dimers stacked onto each other (Maruyama et al., 2013; Mattiroli et al., 2017; Henneman et al., 2018). Thus, it seems that for species in Euryarchaeota, a protein that fulfills the DNA wrapping function (i.e., histone or HTa) seems to be indispensable for the proper folding and regulation of the chromosome. Based on the results of the MNase assay (Figure 3C and 3D), we assume that most crenarchaeal species do not utilize DNA wrapping as the primary level of DNA folding (Figure 7B). Given the efficiency of genome compaction by DNA wrapping, why most crenarchaeal chromosomes do not require DNA wrapping is an interesting question that needs to be answered in the future.

The phylogenetic reconstitution of the HTa/HU gene family has suggested that HTa was acquired through the horizontal transfer of bacterial HU at the root of the clade which includes *Thermoplasmatales* (Hocher et al., 2019). From an evolutionary aspect, an intriguing question is how histone could be replaced by a protein with apparently different characteristics; for example, archaeal histone wraps DNA, whereas bacterial HU bends it (Henneman et al., 2018; Verma et al., 2019). Although some species have been shown to tolerate the complete deletion of histone gene(s) from their genome (Weidenbach et al., 2008), complete deletion of histone genes seems to be impossible in most species (Čuboňová et al., 2012). Besides, since all species in Euryarchaeota (except for *Thermoplasmatales)* encode histone, loss of histone does not seem to be favorable in the living environments. Thus, it seems unlikely that the loss of histone occurred prior to the acquisition of HU. Interestingly, the expression of archaeal histone in *E. coli* does not cause a severe growth defect (Rojec et al., 2019), indicating that histone and HU can coexist in a single organism. These facts are consistent with the idea that the horizontal transfer of bacterial HU to Archaea and its adaptation might have occurred before the loss of the histone gene. It is important to note here that histone is assumed to have existed prior to the acquisition of HU. Without this assumption, other scenarios are possible.

### 3.3 Role of transcription factor-like proteins in *P. calidifontis* and *T. kodakarensis*

Our present results showed the abundance of a putative TF-like protein in the *P. calidifontis* chromosome. Several archaeal TFs and TF-like proteins have been found to combine a global gene regulatory role with an architectural role (Maruyama et al., 2011; Peeters et al., 2015). For example, TrmBL2 (TK0471) in *T. kodakarensis* has a helix-turn-helix motif with non-specific DNA-binding ability (Maruyama et al., 2011; Ahmad et al., 2015). Single-molecule studies have shown that TrmBL2 forms a stiff nucleoprotein filament and that histone competes with TrmBL2, modulating the stiffness of the filament (Efremov et al., 2015). The functions of TrmBL2 resemble those of bacterial H-NS, a NAP that suppresses the transcription of horizontally acquired genetic elements by forming a nucleoprotein filament on genomic DNA (Navarre et al., 2006; Oshima et al., 2006; Higashi et al., 2016). Considering the functional similarity between HNS and TrmBL2 and the fact that HGT occurs between bacteria and Archaea (López-García et al., 2015; Wagner et al., 2017), TrmBL2 might function in suppressing the expression of genes acquired by horizontal transfer in Archaea (Maruyama et al., 2019). Proteins homologous to TrmBL2 have been found in Archaea other than members of the *Thermococcales*, as well as in bacteria (Maruyama et al., 2011). The discovery of Pcal_1183, a TF that belongs to the AsnC/Lrp family, as an abundant chromosomal protein in *P. calidifontis,* adds another candidate HTG suppressor to Archaea. Searching for abundant chromosomal proteins with the method used in the present study would facilitate the identification of novel TF-like nucleoid proteins possibly involved in chromosomal folding or the suppression of HTGs both in bacteria and Archaea in the future.

### 3.4 Conclusions and perspectives

In conclusion, although the chromosomes of Euryarchaeota and Crenarchaeota are commonly folded into higher-order structures, their structural units are fundamentally different. Specifically, DNA wrappers such as histones or HTa are necessary for Euryarchaeota, whereas DNA wrapping as a primary DNA folding is missing in Crenarchaeota (Figure 7A and 7B). Both Euryarchaeota and Crenarchaeota use Alba, which basically functions as a DNA bridger. The function of Alba might be different depending on the interaction with other lineage-specific proteins such as histone or HTa in Euryarchaeota, or Sul7 or Cren7 in Crenarchaeota. TF-like proteins are found in both Euryarchaeota and Crenarchaeota. Possible functions of such TF-like proteins might include architectural roles as well as the suppression of horizontally transferred genetic elements (Maruyama et al., 2019). Further, the interplay between the chromosomal proteins, as well as differential expression and post-translational modifications, might contribute to higher-order folding and regulation of the archaeal chromosomes (Figure 7A).

We also propose that membrane-associated proteins might play roles in chromosome folding and regulation. In the present study, membrane-associated proteins, together with proteins involved in translation, transcription, and replication, were detected by mass spectrometry of proteins associated with isolated archaeal chromosomes. The association between membrane-bound proteins and chromosomes has been proposed in both eukaryotes and bacteria (Maruyama et al., 2014; Chang et al., 2015; Monterroso et al., 2019). In eukaryotes, the linker of nucleoskeleton and cytoskeleton (LINC) complex has been shown to interconnect the chromosome, nuclear membrane, and cytoskeletal filaments, performing diverse functions, including mechanotransduction and meiotic chromosome movements in mice, yeast, and nematodes (Hiraoka and Dernburg, 2009; Sato et al., 2009; Chang et al., 2015). In *E. coli,* the actin-like protein MreB, a member of a membrane-bound complex (MreBCD), is associated with the nucleoid, playing a role in bacterial chromosome segregation (Thanbichler et al., 2005). It can be inferred that archaeal membrane-associated proteins are similarly involved in chromosome segregation. The present findings suggest that membrane-associated proteins might be involved in genome architecture and dynamics in all three domains of life.

## 4. Materials and methods

### 4.1. Strain and growth conditions

*Thermoplasma acidophilum* strain DSM 1728 was grown at 58 °C under aerobic conditions in modified Allan’s basal salt medium (in 1 L: 0.2 g (NH_4_)_2_SO_4_, 3.0 g KH_2_PO_4_, 0.5 g MgSO_4_·7H_2_O, 0.25 g CaCl_2_·_2_ H_2_O) adjusted to pH 1.5 with H_2_SO_4_ and supplemented with 0.1% yeast extract and 1% glucose (Searcy, 1975). *Pyrobaculum calidifontis* strain VA1 was cultivated in the TY medium containing 0.3% sodium thiosulfate pentahydrate at 90 °C (Pelve et al., 2012). *Thermococcus kodakarensis* strain KOD1 was cultured as previously described (Maruyama et al., 2011). *Sulfolobus solfataricus* strain P1 was cultured at 80 °C in *Sulfolobus solfataricus* medium 171 specified by the Japan Collection of Microbes.

### 4.2. Nuclear staining

Live-cell staining of DNA was performed according to a previous study (Poplawski and Bernander, 1997). Briefly, exponentially growing (log) and stationary phase cells were incubated with 1-2 μg/ml of DAPI for 1 hour. These were then harvested, mounted on slides with agarose pads, and visualized using epi-fluorescence microscopy.

### 4.3. On-substrate lysis

On-substrate lysis of archaeal cells was performed as previously described (Maruyama et al., 2011; Ohniwa et al., 2018) with slight modifications. *T. acidophilum* and *P. calidifontis* cells at the log and stationary phases were harvested, washed, and resuspended to a final density of approximately 0.5 OD_600_. The cell suspension was then applied to a round coverslip (15-mm round; Matsunami Glass Ind., Kishiwada, Japan) and dried with nitrogen gas. The cells were lysed with the addition of lysis buffer (10 mM tris pH 6.8, 10 mM EDTA, 0.5% Triton X-100) onto the coverslip and then dried again with nitrogen gas. To achieve sufficient cell lysis, the detergent Triton X-100 was added at the same concentration used for chromosome isolation (see below). The sample was imaged with AFM. To assess the extent of chromosome spreading, samples were stained with DAPI.

### 4.4. Atomic force microscopy

Samples were imaged in the air with the tapping mode of Nanoscope IIIa or IV (Digital Instruments, Santa Barbara, CA) using an OMCL AC160TS probe (Olympus, Tokyo, Japan). Images were captured in a 512 × 512-pixel format, plane-fitted, and flattened using the computer program supplied in the imaging module. Measurements were made, and correction for the “tipeffect” was performed accordingly (Maruyama et al., 2011). Diameters of the chromosomal structures (fibers or particles) were calculated based on the measured bottom-to-bottom distance (apparent measured distance) using the previously described equation S = 0.75W – 16.4, where S is the corrected width of the sample and W is the apparent measured distance (Ohniwa et al., 2007). The contour DNA lengths were measured manually using Nanoscope software; DNA paths were divided into short straight lines and their lengths were summed. When fibers or beadslike structures were encountered, the center of the fiber or beads was used as the DNA path. In this way, the apparent length of DNA wrapped around a protein particle is shortened. When the DNA pathway was not clear, it was excluded from the measurement.

### 4.5. Isolation of archaeal chromosome

Chromosomes from *T. acidophilum*, *P. calidifontis, T. kodakarensis*, and *S. solfataricus* were obtained as described previously with some modifications (Maruyama et al., 2011). Cells were harvested at log or stationary phase and lysed for 10 minutes at 4 °C with extraction buffer (25 mM HEPES, pH 7.0, 15 mM MgCl_2_, 100 mM NaCl, 0.4 M sorbitol, 0.5% Triton X-100). The lysate was then layered on top of a pre-cooled linear sucrose gradient (15–45% sucrose) in 10 mM tris pH 6.8 and 3 mM Mg(CH_3_COO)_2_. The samples were centrifuged in a swinging-bucket MLS-50 rotor (Beckman) at 10,000 × *g* and 4 °C for 20 minutes. The DNA-enriched material, visible as an opalescent band, was collected, washed with MNase buffer (20 mM Tris pH 8.0, 5 mM NaCl, 2.5 mM CaCl_2_), and stored at −20 °C until further use.

### 4.6. MNase digestion of archaeal chromosome

*T. acidophilum, P. calidifontis, T. kodakarensis*, and *S. solfataricus* genomic DNA was incubated with MNase (Worthington Biochemical, Lakewood, NJ) at various concentrations (0.3, 1, 3, 10 and 30 U MNase/100 μl) for 20 minutes at 37 °C. The digestion reaction was stopped by adding stop solution (100 mM EDTA, 100 mM EGTA, 2% SDS). Chromosomal fragments were then purified with phenol/chloroform extraction and ethanol precipitation. The DNA pellet was dissolved in TE buffer (10 mM tris pH 7.5, 1 mM EDTA), resolved with a 2.5% agarose gel, and visualized with ethidium bromide staining.

### 4.7. Mass spectrometry

Chromosomes were isolated from cells in the stationary phase as described previously herein, digested with MNase (Worthington Biochemical Corp., Lakewood, NJ), and run on a 15% SDS-PAGE gel to separate proteins associated with archaeal chromosomes. After staining with Coomassie Brilliant Blue, visible protein bands were excised from the gel and processed for liquid chromatography coupled with tandem mass spectrometry (LC-MS/MS) as described previously (Maruyama et al., 2011). Top hits of each band are summarized in Tables 1–3.

**Table 1.**
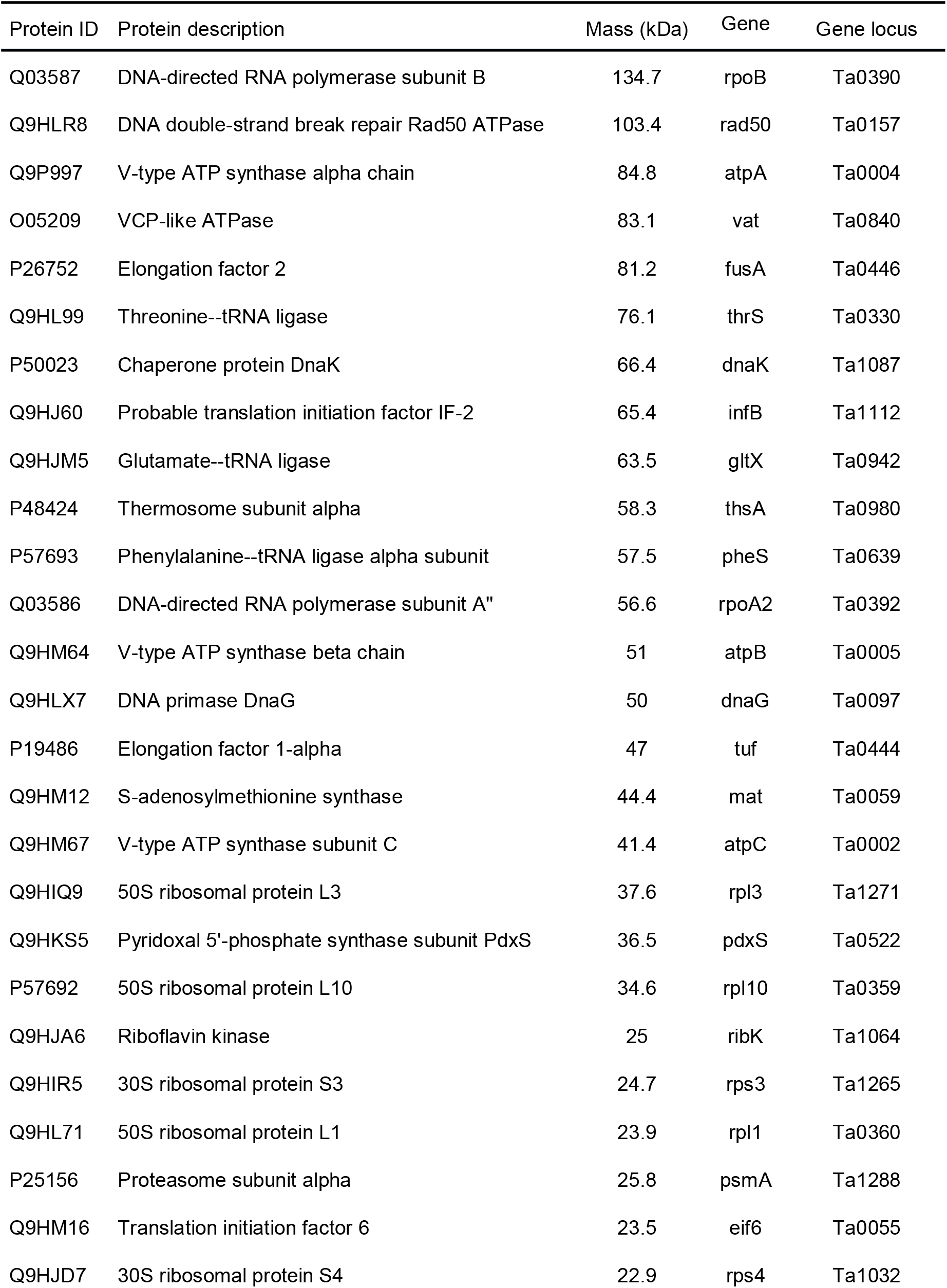
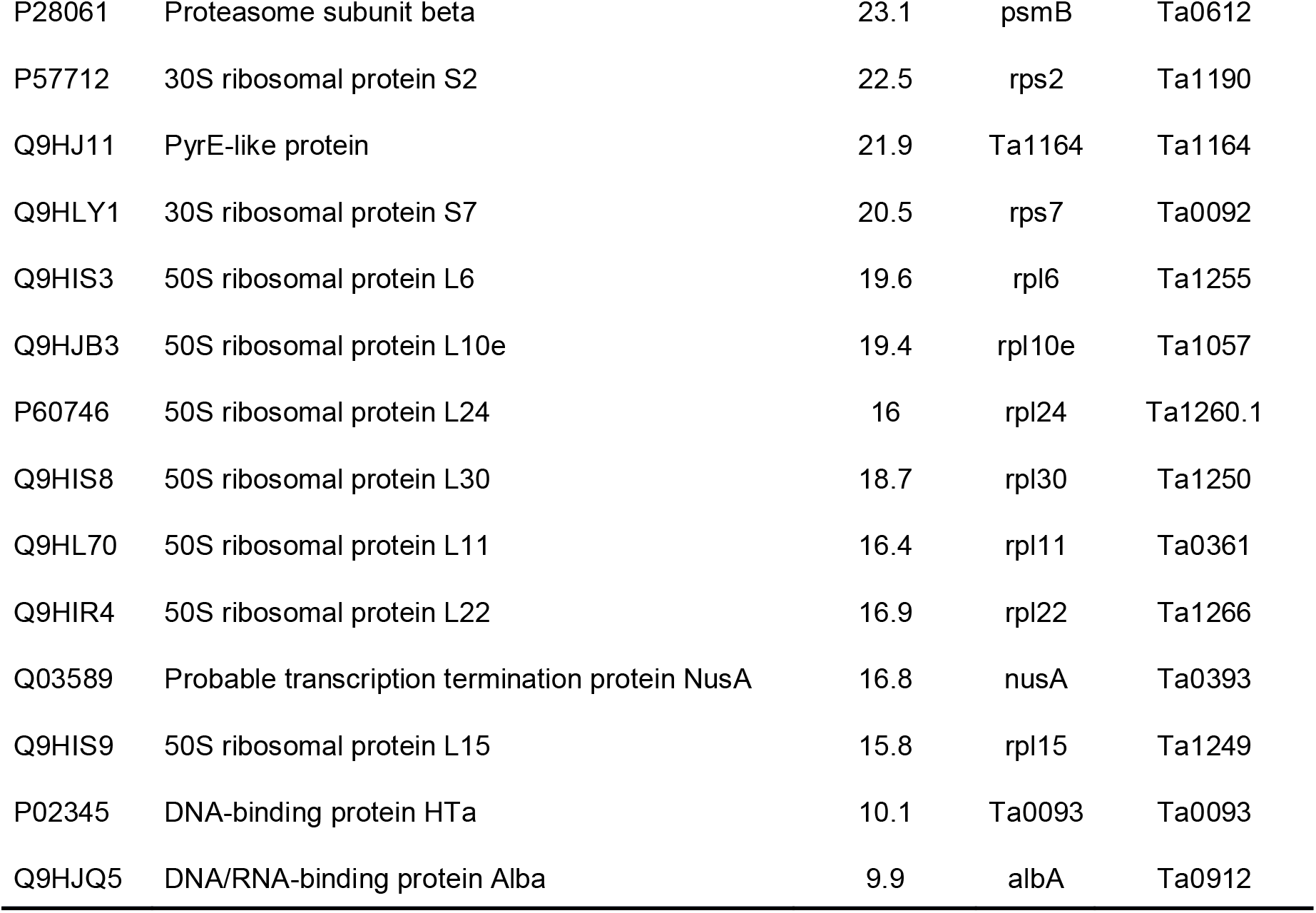
Proteins identified by mass spectrometry from isolated chromosomes of *Thermoplasma acidophilum*. Protein and gene information based on the UniProt database (The UniProt Consortium, 2019).

**Table 2.** Proteins identified by mass spectrometry from isolated chromosomes of *Pyrobaculum calidifontis*

| Protein | Protein description | Mass | Gene name | Gene locus |
| --- | --- | --- | --- | --- |
| A3MWL6 | Pullulanase / alpha-amylase | 111.5 | Pcal_1616 | Pcal_1616 |
| A3MUT3 | Oligopeptide binding protein, putative | 95.6 | Pcal_0975 | Pcal_0975 |
| A3MTK9 | V-type ATP synthase subunit I | 85.8 | Pcal_0549 | Pcal_0549 |
| A3MY13 | AAA family ATPase, CDC48 subfamily | 81.8 | Pcal_2115 | Pcal_2115 |
| A3MY25 | Succinate dehydrogenase subunit A | 64.5 | Pcal_2127 | Pcal_2127 |
| A3MV10 | Thermosome subunit | 60.1 | Pcal_1052 | Pcal_1052 |
| A3MX21 | Thermosome subunit | 60.2 | Pcal_1771 | Pcal_1771 |
| A3MV59 | ATPase, PilT family | 58 | Pcal_1101 | Pcal_1101 |
| A3MV69 | Elongation factor 1-alpha | 48.9 | tuf | Pcal_1111 |
| A3MV64 | rRNA biogenesis protein Nop56/Nop58 | 46.8 | Pcal_1106 | Pcal_1106 |
| A3MV08 | DNA primase DnaG | 45 | dnaG | Pcal_1050 |
| A3MY01 | S-adenosylmethionine synthase | 44.4 | mat | Pcal_2103 |
| A3MX84 | 50S ribosomal protein L10 | 37.5 | rpl10 | Pcal_1834 |
| A3MUU8 | DNA polymerase sliding clamp | 29.4 | pcn | Pcal_0990 |
| A3MY29 | DNA polymerase sliding clamp | 28.5 | pcn | Pcal_2131 |
| A3MWF6 | Thiamine thiazole synthase | 27.7 | thi4 | Pcal_1555 |
| A3MX83 | 50S ribosomal protein L1 | 25.1 | rpl1 | Pcal_1833 |
| A3MU88 | 30S ribosomal protein S5 | 22.1 | rps5 | Pcal_0779 |
| A3MXZ7 | 50S ribosomal protein L13 | 21.7 | rpl13 | Pcal_2099 |
| A3MTM7 | DJ-1_Pfpl domain-containing protein | 20.9 | Pcal_0567 | Pcal_0567 |
| A3MXA2 | Uncharacterized protein | 20 | Pcal_1852 | Pcal_1852 |
| A3MVE1 | Transcriptional regulator, AsnC family | 18.7 | Pcal_1183 | Pcal_1183 |
| A3MUQ5 | DNA/RNA-binding protein Alba | 9.9 | alba | Pcal_0947 |

**Table 3.** Proteins identified by mass spectrometry from isolated chromosomes of *Thermococcus kodakarensis*

| Protein ID | Protein description | Mass (kDa) | Gene | Gene locus |
| --- | --- | --- | --- | --- |
| Q5JH54 | Predicted endonuclease-methyltransferase fusion protein | 150.7 | TK1460 | TK1460 |
| Q5JE98 | CDC48/VCP homolog, AAA superfamily | 88.9 | TK1157 | TK1157 |
| Q5JE33 | DNA-directed RNA polymerase subunit | 102.9 | TK1082 | TK1082 |
| Q5JH85 | Putative 5-methylcytosine restriction system, GTPase subunit | 79.7 | TK0795 | TK0795 |
| Q5JHC8 | Acylamino acid-releasing enzyme | 72.1 | TK0752 | TK0752 |
| Q5JH81 | Type 2 DNA topoisomerase 6 subunit B | 64.2 | top6B | TK0799 |
| Q5JCX3 | AMP phosphorylase | 53.7 | deoA | TK0352 |
| Q5JIR2 | V-type ATP synthase beta chain | 52.4 | atpB | TK1603 |
| Q5JFZ4 | Elongation factor 1-alpha | 47.5 | tuf | TK0308 |
| Q5JE34 | DNA-directed RNA polymerase subunit A'' | 43.7 | rpoA2 | TK1081 |
| Q5JGA2 | Uncharacterized protein | 40.8 | TK0462 | TK0462 |
| Q5JDJ0 | 50S ribosomal protein L3 | 39 | rpl3 | TK1542 |
| Q5JHG6 | Uncharacterized protein | 36.7 | TK2157 | TK2157 |
| Q5JHT2 | Transcription regulator, SpoVT/AbrB family, containing PhoU domain | 35.3 | TK2245 | TK2245 |
| Q5JD67 | HTH-type transcriptional regulator TrmBL2 | 30.8 | trmBL2 | TK0471 |
| Q5JGL5 | Uncharacterized protein | 29.3 | TK1263 | TK1263 |
| Q5JFN1 | Fibrillarin-like rRNA/tRNA 2'-O-methyltransferase | 25.3 | flpA | TK0183 |
| Q5JF30 | Peroxiredoxin | 24.6 | TK0537 | TK0537 |
| Q5JDD5 | Transcription factor E | 22.1 | tfe | TK2024 |
| Q5JG73 | Iron-molybdenum cofactor-binding protein | 21.5 | TK0724 | TK0724 |
| Q5JJH0 | 50S ribosomal protein L15 | 16.5 | rpl15 | TK1519 |
| Q5JH90 | UPF0179 protein TK0790 | 15.9 | TK0790 | TK0790 |
| Q5JJF8 | 50S ribosomal protein L14 | 15.2 | rpl14 | TK1531 |
| Q5JF39 | DNA/RNA-binding protein Alba | 10.1 | albA | TK0560 |
| Q9Y8I1 | Archaeal histone A | 7.4 | hpkA | TK1413 |
| Q9Y8I2 | Archaeal histone B | 7.2 | hpkB | TK2289 |

### 4.8. Expression and purification of recombinant proteins

Genes encoding the protein Alba from *T. acidophilum* and *P. calidifontis* and HTa from *T. acidophilum* were PCR-amplified from their respective genomic DNA template and inserted into a pET16b vector containing 10x histidine tag at the N-terminus (Novagen, Madison, WI). The following primers were used: for *T. acidophilum* Alba, forward 5’-ATCGCCATGGCAGAGGAGAACATAATCTTTG-3’ and reverse 5’-ATCGGGATCCTCAACGTGACAGC-3’; for *P. calidifontis* Alba, forward 5’-ATCGCCATGGCGACAGAACAGACAATAC-3’ and reverse 5’-ATCGGATCCTTAGGCCATTTCGAGCACT-3’; for *T. acidophilum* HTa, forward 5’-ATCGCATATGTAGGAATCAGTGAGCTAT-3’ and reverse 5’-ATCGGGATCCTTACTGCTGGTATTTTATCTTGC −3’. The Alba gene was inserted using the NcoI and BamHI restriction sites, resulting in the removal of the his-tag. HTa gene was inserted via the NdeI and BamHI sites and thus tagged with histidine. Competent *E. coli* BL21-CodonPlus (DE3)-RIL was transformed with the resulting constructs. Protein expression was induced via the addition of IPTG. Recombinant proteins were then extracted and purified. Alba proteins were run on an ion-exchange column (HiTrap SP column, 1 ml, GE Healthcare UK Ltd. Buckinghamshire, England) following the manufacturer’s protocol and as described previously (Maruyama et al., 2011). The His-tagged HTa was purified with Ni-NTA agarose beads (Qiagen) following the manufacturer’s protocol. Histones from *T. kodakarensis* were expressed and purified as previously described (Maruyama et al., 2011).

### 4.9. In vitro reconstitution

The plasmid pBlueScriptII (Agilent Technologies Inc., Santa Clara, CA) was digested with Hind III restriction enzyme digestion, resulting in a 2961-bp linearized double-stranded DNA. Each recombinant protein was mixed with the 3-kb linearized DNA at various DNA-to-protein weight ratios in 10 mM tris pH 6.8 and 200 mM NaCl. In each experiment, DNA and the protein were mixed in a 20 μl volume. The DNA concentration was fixed to 5 ng/μl (100 ng total). Protein concentration was included at a concentration that achieves the desired DNA:protein weight ratio. The mixture was heated for 10 minutes at appropriate temperatures, 58 °C for *T. acidophilum* and 90 °C for *P. calidifontis*, incubated for 10 minutes at 25 °C, and then processed for AFM imaging. For AFM observation, the samples were diluted 1:20 with AFM fixation buffer (10 mM tris pH 8.0, 5 mM NaCl, 0.3% glutaraldehyde) and incubated for 30 minutes at 25 °C. The protein-DNA complexes were then deposited onto a freshly cleaved mica surface pretreated with 10 mM spermidine. After 10 minutes, the mica was washed with 1 ml of pure water, dried with nitrogen gas, and observed with AFM. Note that the theoretical length of a 2961-bp dsDNA in B-form is ~1007 nm (0.34 nm/bp). The theoretical length is indicated in the histograms as a dashed line. Reconstitution experiments using a combination of different chromosomal proteins were also performed by mixing the 3-kb linearized DNA with the protein pair of interest at specified amounts. In this case, the protein weight ratio are indicated relative to DNA (e.g. 2:6 HTa:Alba means 1:2:6 DNA:HTa:Alba weight ratio).

## 7. Acknowledgments

We would like to thank Mr. Yuzo Watanabe (Proteomics Facility, Graduate School of Biostudies, Kyoto University) and Dr. Ryosuke Ohniwa (Faculty of Medicine, University of Tsukuba) for mass spectrometry assistance and Editage (www.editage.com) for English editing.

## 8. Author contributions

HM and KT conceived and designed the study. EP performed experiments, analyzed data, and wrote the first draft of the manuscript. TN, CM, and TO wrote sections of and gave critical comments on the manuscript. HM, EP, HA, and KT revised the manuscript.

